# Complete plastome sequences from *Bertholletia excelsa* and 23 related species yield informative markers for Lecythidaceae

**DOI:** 10.1101/192112

**Authors:** Ashley M. Thomson, Oscar M. Vargas, Christopher W. Dick

## Abstract

- *Premise of the study:* The tropical tree family Lecythidaceae has enormous ecological and economic importance in the Amazon basin. Lecythidaceae species can be difficult to identify without molecular data, however, and phylogenetic relationships within and among the most diverse genera are poorly resolved.
- *Methods:* To develop informative genetic markers for Lecythidaceae, we used genome skimming to assemble *de novo* the full plastome of the Brazil nut tree *(Bertholletia excelsa)* and 23 other Lecythidaceae species. Indices of nucleotide diversity and phylogenetic signal were used to identify regions suitable for genetic marker development.
- *Results:* The *B. excelsa* plastome contained 160,472 bp and was arranged in a quadripartite structure. Using the 24 plastome alignments, we developed primers for 10 coding and non-coding DNA regions containing exceptional nucleotide diversity and phylogenetic signal. We additionally developed 19 chloroplast simple sequence repeats (cpSSRs) for population-level studies.
- *Discussion:* The coding region *ycf1* and the spacer *rpl16-rps3* outperformed plastid DNA markers previously used for barcoding and phylogenetics. Used in a phylogenetic analysis, the matrix of 24 plastomes showed with 100% bootstrap support that *Lecythis* and *Eschweilera* are polyphyletic. The plastomes and primers presented in this study will facilitate a broad array of ecological and evolutionary studies in Lecythidaceae.

## INTRODUCTION

Lecythidaceae *(sensu latu)* is a pantropical family of trees with three subfamilies: Foetidioideae, which is restricted to Madagascar; Planchonioideae, found in the tropical forests of Asia and Africa; and the Neotropical clade Lecythidoideae (Mori et al., 2007), which contains ca. 234 of the ca. 278 known species in the broader family (Mori et al., 2007; Huang et al., 2015; Mori et al., 2017; Mori, 2017). Neotropical Lecythidaceae are understory, canopy, or emergent trees with distinctive floral morphology and woody fruit capsules. Among Lecythidaceae species are the iconic Brazil nut tree, *Bertholletia excelsa* Bonpl.; the oldest documented angiosperm tree, *Cariniana micrantha* Ducke (dated at >1400 years old in Manaus, Brazil; Chambers et al., 1998); the cauliflorous cannonball tree commonly grown in botanical gardens, *Couroupita guianensis* Aubl.; and important timber species (e.g. *Carinaria legalis* (Mart.) Kuntze). Lecythidaceae is the third most abundant family of trees in the Amazon forest, following Fabaceae and Sapotaceae (ter Steege et al., 2013). The most species-rich genus, *Eschweilera*, with ca. 99 species (Mori, 2017), is also the most abundant tree genus in the Amazon basin (ter Steege et al., 2013), and *Eschweilera coriacea* (DC.) S.A.Mori is the most common tree species in much of Amazonia (ter Steege et al., 2013). Lecythidaceae provide important ecological services such as carbon sequestration, and food resources for pollinators (bats and large bees) and seed dispersers (monkeys and agouties) (Prance and Mori, 1979; Mori and Prance, 1990).

Tools for species-level identification and phylogenetic analyses of Lecythidaceae could significantly advance research on Amazon tree diversity. However, despite their ease of identification at the family level, species-level identification of many Lecythidaceae (especially *Eschweilera)* is notoriously difficult when based on sterile (i.e. without fruit or floral material) herbarium specimens, and flowering of individual trees often occurs only at multi-year intervals (Mori and Prance, 1987). As a complement to other approaches, DNA barcoding (Dick and Kress, 2009; Dexter et al., 2010) may help to identify species or clades of Lecythidaceae.

A combination of two protein-coding plastid regions *(matK* and *rbcL)* have been proposed as core plant DNA barcodes (Hollingsworth et al., 2009), although other coding and non-coding plastome regions *(psbA-trnH, rpoB, rpoC1, trnL, ycf5)* and the internal transcribed spacer (ITS) of nuclear ribosomal genes, have been recommended as supplemental barcodes for vascular plants (Kress et al., 2005; Lahaye et al., 2008; Li et al., 2011). However, an evaluation of a subset of these markers (ITS, *psbA-trnH, matK, rbcL, rpoB, rpoC1, trnL)* on Lecythidaceae in French Guiana (Gonzales et al., 2009) showed poor performance for species identification. Furthermore, the use of traditional markers (plastid *ndhF, trnL-F*, and *trnH-psbA*, and nuclear ITS) for phylogenetic analysis has produced weakly supported trees (Mori et al., 2007; Huang et al., 2015) indicating a need to develop more informative markers and/or increase molecular sampling.

The main objectives of this study were to (1) assemble, annotate, and characterize the first complete plastome sequence of Lecythidaceae, from the iconic Brazil nut tree *Bertholletia excelsa;* (2) obtain a robust backbone phylogeny for the Neotropical clade using newly-assembled draft plastome sequences for an additional 23 species; and (3) develop a novel set of informative molecular markers for DNA barcoding and broader evolutionary studies.

## METHODS

**Plant material and DNA library preparation—**We performed genomic skimming on 24 Lecythidaceae species, including 23 Lecythidoideae and one outgroup species *(Barringtonia edulis* Seem.) from the Planchonioideae. The sampling included all 10 Lecythidoideae genera (Appendix 1). Silica-dried leaf tissue from herbarium-vouchered collections was collected by Scott Mori and colleagues and loaned by the New York Botanical Garden. Total genomic DNA was extracted from 20 milligrams of dried leaf tissue using the NucleoSpin Plant II extraction kit (Machery-Nagel, Bethlehem, PA, USA) with SDS lysis buffer. Prior to DNA library preparation, 5 micrograms of total DNA were fragmented using a Covaris S-series sonicator (Covaris, Inc. Woburn, MA, USA), following the manufacturer’s protocol, to obtain ca. 300 bp insert-sizes. We prepared the sequencing library using the NEBNext DNA library Prep Master Mix and Multiplex Oligos for Illumina Sets (New England BioLabs Inc. Ipswich, MA, USA) according to the manufacturer’s protocol. Size-selection was carried out prior to PCR using Pippin Prep (Sage Science, Beverly, MA, USA). Molecular mass of the finished paired-end library was quantified using an Agilent 2100 Bioanalyzer (Agilent Technologies Inc., Santa Clara, CA, USA) and by qPCR using an ABI PRISM 7900HT (ThermoFisher Scientific, Waltham, MA, USA) at the University of Michigan DNA Sequencing Core (Ann Arbor, MI, USA). We sequenced the libraries on one lane of the Illumina HiSeq 2000 (Illumina Inc., San Diego, CA, USA) with a paired-read length of 100 bp.

***Plastome assembly*—**Illumina adaptors and barcodes were excised from raw reads using Cutadapt v.1.4.2 (Martin, 2011). Reads were then quality-filtered using Prinseq v. 0.20.4 (Schmieder and Edwards, 2011), which trimmed 5’ and 3’ sequence ends with Phred quality score <20 and removed all trimmed sequences <50 bp in length, with >5% ambiguous bases, or with mean Phred quality score <20. A combination of *de novo* and reference-guided approaches were used to assemble the plastomes. First, chloroplast reads were separated from the raw read pool by Blast-searching all raw reads against a database consisting of all complete angiosperm plastome sequences available on GenBank (accessed in 2014). Any aligned reads with an e-value <1^5^ were retained for subsequent analysis. The filtered chloroplast reads were *de novo* assembled using Velvet v.7.0.4 (Zerbino and Birney, 2008) with kmer values of 71, 81, and 91 using a low-coverage cutoff of 5 and minimum contig length of 300. The assembled contigs were then mapped to a reference genome (see below) using Geneious v. R8 (Kearse et al., 2012) to determine their order and direction using the reference-guided assembly tool with medium sensitivity and iterative fine-tuning options. Finally, raw reads were iteratively mapped onto the draft genome assembly to extend contigs and fill gaps using low-sensitivity reference-guided assembly in Geneious. We first assembled the draft genome of *Bertholletia excelsa;* the plastomes of the remaining 23 species were assembled subsequently using the plastome of *B. excelsa* as a reference. The *B. excelsa* plastome was annotated using DOGMA (Wyman et al., 2004) with the default settings for chloroplast genomes. Codon start and stop positions were determined using the open reading frame finder in Geneious and by comparison with the plastome sequence of *Camellia sinensis* var. *pubilimba* Hung T. Chang (Genbank ID: KJ806280). A circular representation of the *B. excelsa* plastome was made using OGDraw V1.2 (Lohse et al., 2007). The complete annotated plastome of *B. excelsa* and the draft plastomes of the remaining 23 Lecythidaceae species sampled were deposited in GenBank (Appendix 1).

***Identification of molecular markers*—**Chloroplast simple sequence repeats (cpSSRs) in *B. excelsa* were identified using the Phobos Tandem Repeat Finder v.3.3.12 (Mayer, 2010) by searching for uninterrupted repeats of nucleotide units of 1 to 6 bp in length, with thresholds of ≥12 mononucleotide, ≥6 dinucleotide, and ≥4 trinucleotide, and ≥3 tetra-, penta-, and hexanucleotide repeats (Sablok et al., 2015). We developed primers to amplify the cpSSRs using Primer 3 v.2.3.4 (Untergasser et al., 2012) with the default options and setting the PCR product size range between 100 and 300 bp.

The 24 plastomes were aligned with MAFFT v.7.017 (Katoh et al., 2002) and scanned for regions of high nucleotide diversity, π (Nei 1987), using a sliding window analysis implemented in DNAsp v.5.10.1 (Librado and Rozas, 2009) with a window and a step size of 600 bp. Levels of nucleotide diversity were plotted using the generic function “plot” in R (R core development group), and windows with values over the 95^th^ percentile were considered of high π.

Taking into account that DNA barcodes can also be used in phylogenetic analyses and because regions with high π do not necessarily have high phylogenetic signal (e.g. unalignable hypervariable regions), we employed a log-likelihood approach modified from Walker et al. (2017) to identify phylogenetically influential regions. First, we inferred a phylogenetic tree with the plastome alignment (including only one inverted repeat) by performing 100 independent maximum likelihood (ML) searches using a GTRGAMMA model with RAxML v. 8.2.9 (Stamatakis, 2014). Those searches resulted in the same topology that was subsequently annotated with the summary from 100 bootstraps using “sumtrees.py" v.4.10 (Sukumaran and Holder, 2010). Then, we calculated the site-specific log-likelihood in the alignment over the plastome phylogeny and calculated their differences site-wise to the averaged log-likelihood per site of 1000 randomly permuted trees (tips were randomly shuffled). Log-likelihood scores were calculated with RAxML under using a GTRGAMMA model. The site-wise log-likelihood differences (LD) were calculated using 600 bp non-overlapping windows with a custom R script (see below). We interpreted greater LD as an indication of greater phylogenetic signal, and windows with LD above the 95^th^ percentile were considered to have exceptional phylogenetic signal.

Primers flanking the top ten regions with high π were designed using Primer 3 with default program options. We employed a maximum product size of 1300 bp because lower cutoffs values (i.g. 600 bp) made the primer design extremely challenging due to the lack of conserved regions. Primers were designed to amplify across all 23 Neotropical species without the use of degenerate bases. However, primers with a small number of degenerate bases were permitted for some regions where primer development otherwise would not have been possible due to high sequence variability in the priming sites. We investigated the potential of our markers to produce robust phylogenies by calculating individual gene trees in RAxML v.8.2.9 in an ML search with 100 rapid bootstraps (option “-f a”) using the GTRGAMMA model. To evaluate the number of markers needed to obtain a resolved tree with an average of ~90 bootstrap support (BS), we first concatenated the two markers with the highest π and inferred a tree; subsequently we added the marker with the next highest π score. We iterated this process until we obtained a matrix with each of the 10 markers developed. For every tree obtained, we calculated its average BS and its Robinson-Foulds distance (RF) (Robinson and Foulds, 1981) from the plastome phylogeny, using a custom R script employing the packages APE (Paradis et al., 2004) and Phangor (Schliep 2011). Scripts and alignments used for this study can be found at https://bitbucket.org/oscarvargash/lecythidaceae_plastomes.

## RESULTS

***Lecythidaceae plastome features*—**The sequenced plastome of *Bertholletia excelsa* contained 160,472 base pairs and 115 genes, of which 4 were rRNAs and 30 were tRNAs (Fig. 1, Table 1). The arrangement of the *B. excelsa* plastome had a typical angiosperm quadripartite structure with a single copy region of 85,830 bp, a small single copy region of 16,670 bp, and two inverted of repeats of 27,481 bp each. Relative to *Camellia sinensis* var. *pubilimba*, we found no gene gain/losses in *B. excelsa*. The only structural difference found is that *B. excelsa* contains the sequential genes *trnH-GUG, rps3, rpl22*, and *rps19* in the inverted repeat while *C. sinensis* var. *pubilimba* contains these genes in the large single copy region. Similarly, no gene gain/losses are found when *B. excelsa* is compared to other neotropical Lecythidaceae plastomes assembled herein (Table 2). In addition to *B. excelsa*, the plastome of *Eschweilera alata* was also completely assembled; the coverage for the remaining plastomes ranged between 85% and 99.60% (Appendix 1).

**Fig. 1.**
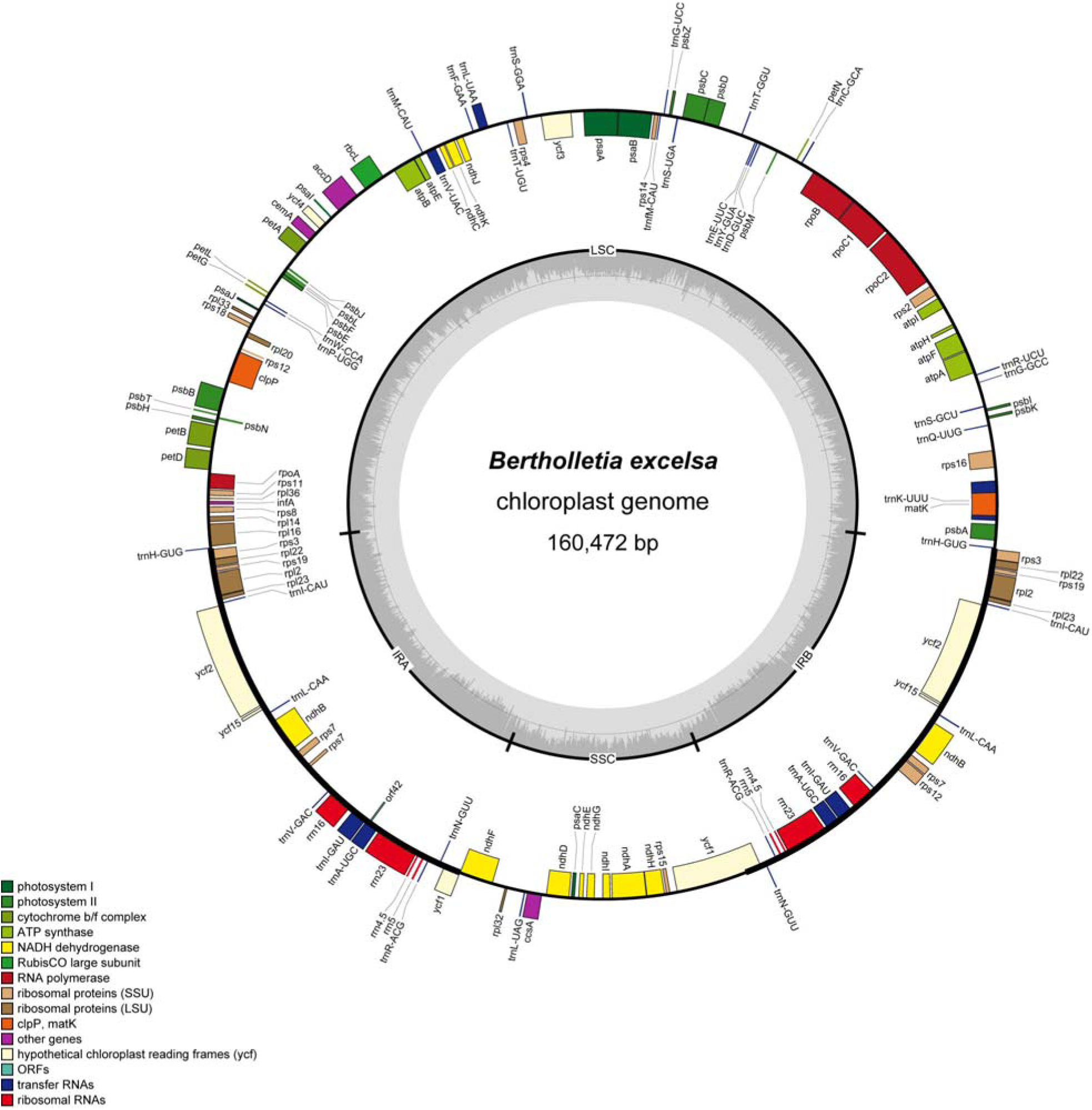
Plastome map of the Brazil-nut tree *Bertholletia excelsa.* Genes outside the circle are transcribed clockwise, genes inside the circle are transcribed counter-clockwise. Gray bars in the inner ring show the GC content percentage.

**Table 1.**
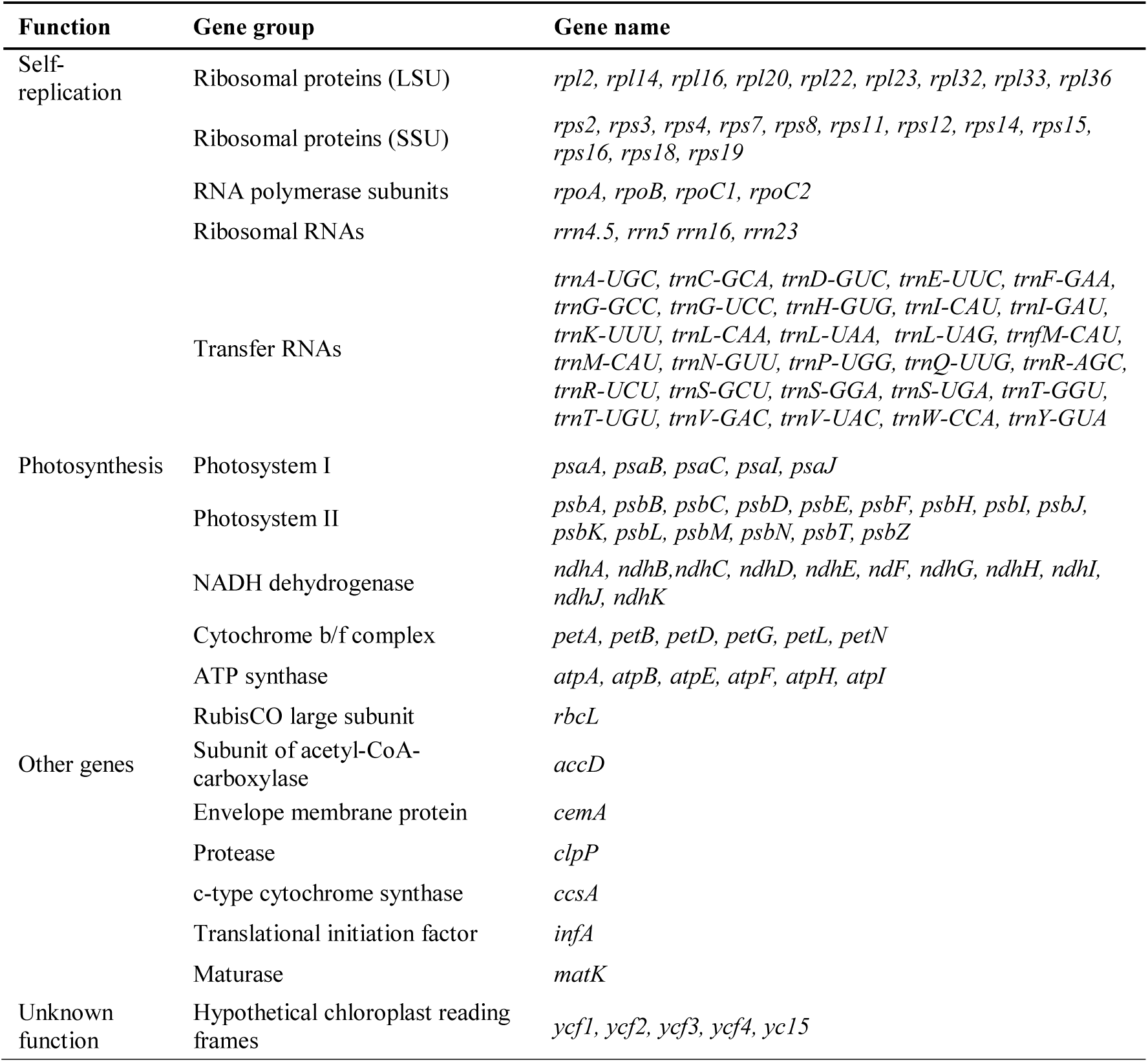
Genes contained within the chloroplast genome of *Bertholletia excelsa.*

**Table 2.**
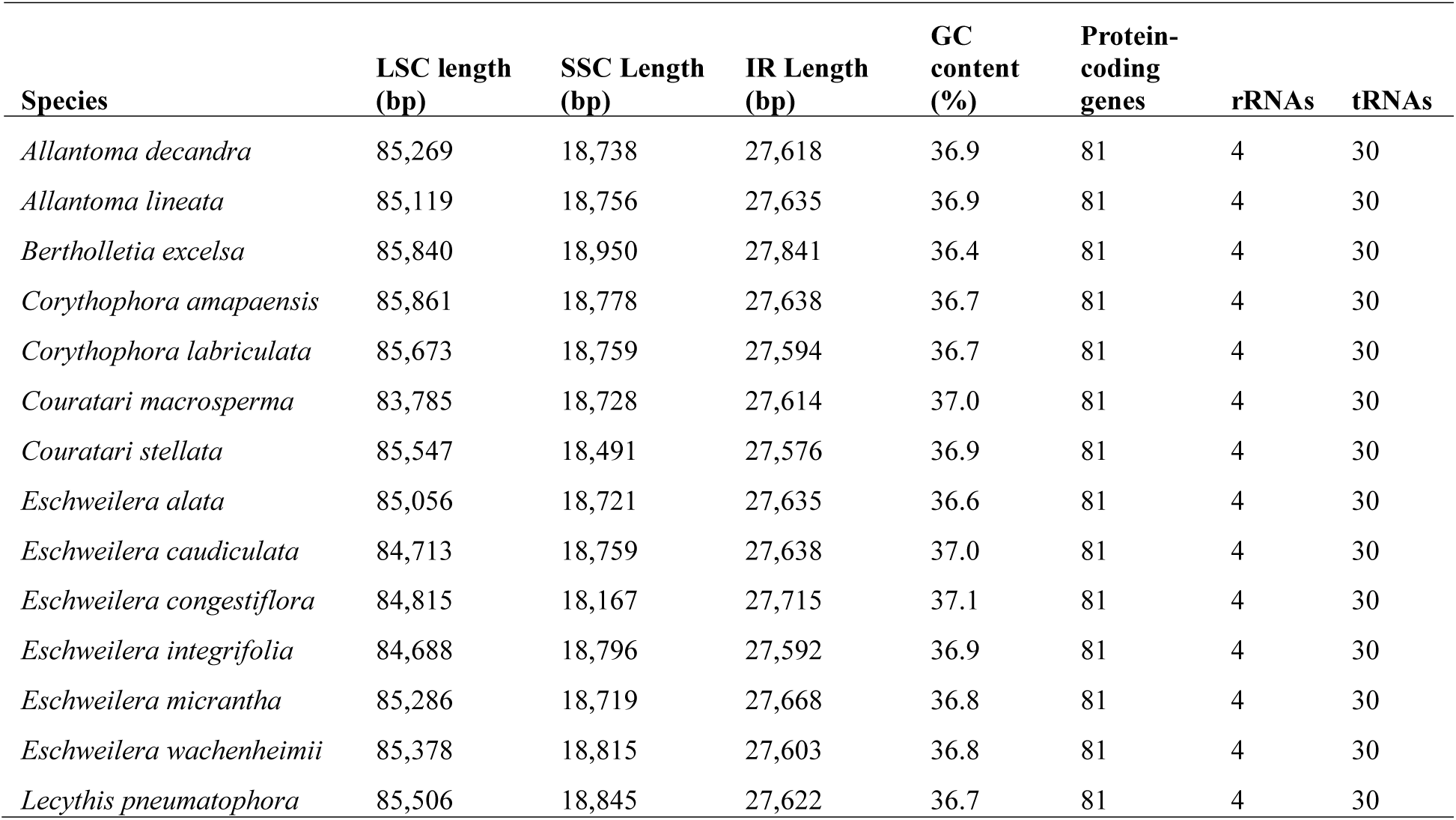
Comparison for plastome subunits for the samples for which the inverted repeats (IR) were completely assembled. Length and GC content of the large single copy (LSC) and small single copy (SSC) regions in partial plastomes are estimates only.

| Species | LSC length<br>(bp) | SSC Length<br>(bp) | IR Length<br>(bp) | GC<br>content<br>(%) | Protein-coding<br>genes | rRNAs | tRNAs |
| --- | --- | --- | --- | --- | --- | --- | --- |
| *Allantoma decandra* | 85,269 | 18,738 | 27,618 | 36.9 | 81 | 4 | 30 |
| *Allantoma lineata* | 85,119 | 18,756 | 27,635 | 36.9 | 81 | 4 | 30 |
| *Bertholletia excelsa* | 85,840 | 18,950 | 27,841 | 36.4 | 81 | 4 | 30 |
| *Corythophora amapaensis* | 85,861 | 18,778 | 27,638 | 36.7 | 81 | 4 | 30 |
| *Corythophora labriculata* | 85,673 | 18,759 | 27,594 | 36.7 | 81 | 4 | 30 |
| *Couratari macrosperma* | 83,785 | 18,728 | 27,614 | 37.0 | 81 | 4 | 30 |
| *Couratari stellata* | 85,547 | 18,491 | 27,576 | 36.9 | 81 | 4 | 30 |
| *Eschweilera alata* | 85,056 | 18,721 | 27,635 | 36.6 | 81 | 4 | 30 |
| *Eschweilera caudiculata* | 84,713 | 18,759 | 27,638 | 37.0 | 81 | 4 | 30 |
| *Eschweilera congestiflora* | 84,815 | 18,167 | 27,715 | 37.1 | 81 | 4 | 30 |
| *Eschweilera integrifolia* | 84,688 | 18,796 | 27,592 | 36.9 | 81 | 4 | 30 |
| *Eschweilera micrantha* | 85,286 | 18,719 | 27,668 | 36.8 | 81 | 4 | 30 |
| *Eschweilera wachenheimii* | 85,378 | 18,815 | 27,603 | 36.8 | 81 | 4 | 30 |
| *Lecythis pneumatophora* | 85,506 | 18,845 | 27,622 | 36.7 | 81 | 4 | 30 |

***Identification of molecular markers*—**Within the plastome of *Bertholletia excelsa* we found 23 cpSSRs, 22 of which were in non-coding regions and one in the *ndhD* coding region. We designed 19 primer pairs with an acceptable product length, annealing temperature, and GC content for cpSSRs located in noncoding regions (Table 3). π exceeded the 95^th^ percentile for nine 600 bp windows (Fig. 2A, Table 4, 5). Similarly, 13 windows were over the 95^th^ percentile for LD (Fig. 2B, Table 4, 5) indicating high phylogenetic signal. While most of the informative windows were in non-coding regions, two consecutive regions were positioned in the *ycf1* gene. Six windows contained both high π and LD. As expected, high π and greater LD largely agreed. Based on the rank of the windows obtained for π, we developed primers for the following regions (ordered from high to low π): *ycf1, rpl16-rps3, psbM-trnD, ccsA-ndhD, trnG-psaB, petD-rpoA, psbZ-trnfM, trnE-trnT*, and *trnT-psbD* (Table 6).

**Table 3.**
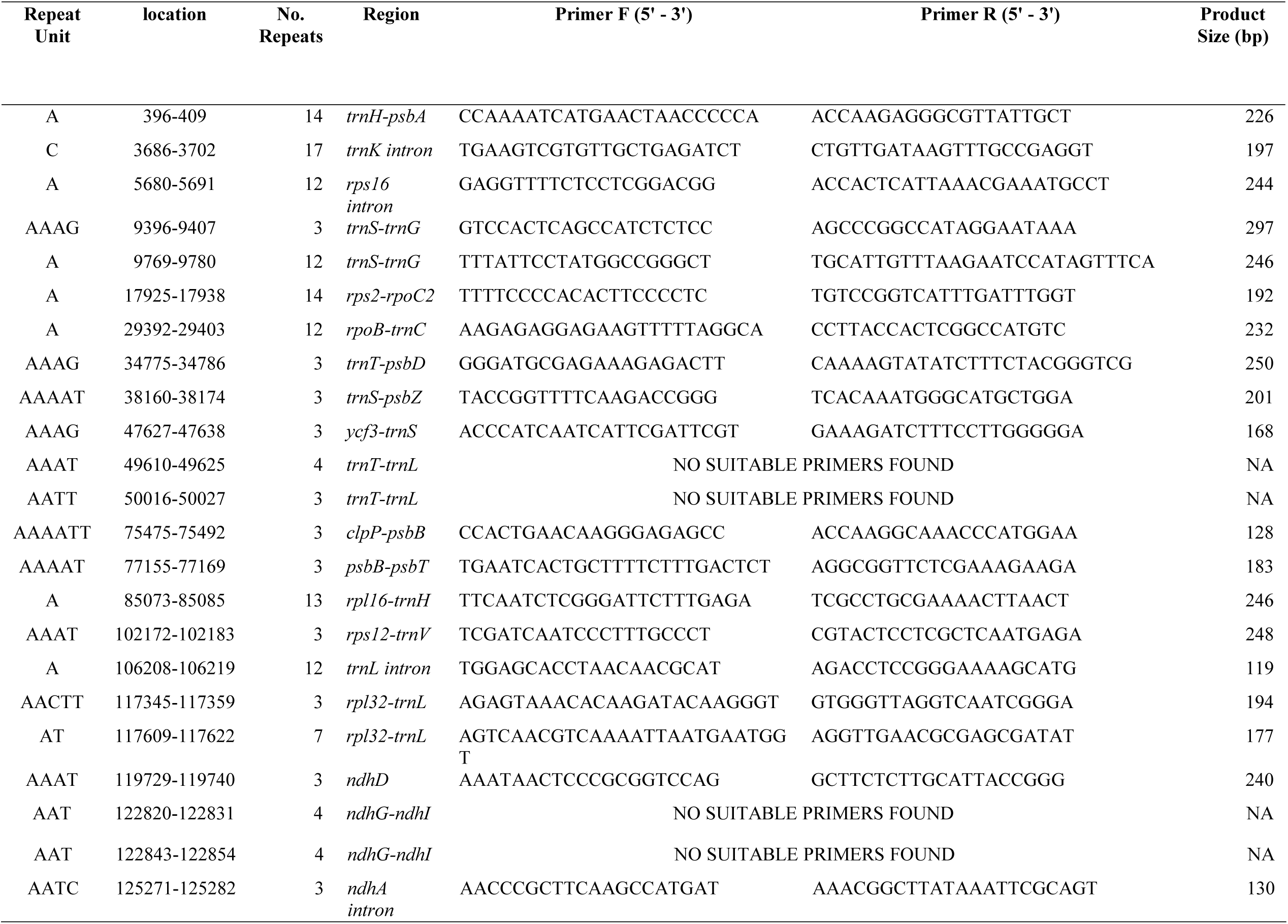
Primers for the amplification of simple sequence repeats in the plastome of *Bertholletia excelsa.* All primers pairs amplify non-coding sequences with the exception of *ndhD.*

**Table 4.** Regions of the chloroplast binned in windows of 600 sites with high (above the 95^th^ percentile) nucleotide diversity (π) and/or site-wise log-likelihood score differences (LD). LSC: large single copy. SSC: small single copy (see main text). Coding regions are indicated in windows that have the same 5’ and 3’ expressed flanking regions in column 3. Notice that no regions are reported for the inverted repeat (IR). Coordinates are given on the alignment and the *Bertholletia excelsa* plastome that are assembled with the standard LSC-SSC-IR structure.

| Location in the alignment | *Bertholletia* plastome location | Closest flanking expressed region | | Region | $\pi$ | LD |
| --- | --- | --- | --- | --- | --- | --- |
|  |  | 5' | 3' |  |  |  |
| 1–600 | 1–490 | *trnH* | *psbA* | LSC |  | * |
| 5401–6000 | 4885–5373 | *trnK-UUU* | *rps16* | LSC |  | * |
| 34801–35400 | 30925–31450 | *petN* | *trnD-GUC* | LSC |  | * |
| 35401–36000 | 31451–31967 | *psbM* | *trnD-GUC* | LSC | * | * |
| 37201–37800 | 33027–33573 | *trnE-UUC* | *trnT-GGU* | LSC | * | * |
| 39601–40200 | 34893–35433 | *trnT-GGU* | *psbD* | LSC |  | * |
| 43801–44400 | 38798–39254 | *psbZ* | *trnfM-CAU* | LSC | * | * |
| 44401–45000 | 39255–39744 | *trnfM-CAU* | *psaB* | LSC | * | * |
| 61201–61800 | 54771–55275 | *trnV-UAC* | *atpE* | LSC |  | * |
| 78601–79200 | 70230–70771 | *psaJ* | *rps18* | LSC |  | * |
| 89401–90000 | 80536–81103 | *petD* | *rpoA* | LSC | * |  |
| 95401–96000 | 85455–85906 | *rpl16* | *rps3* | LSC | * |  |
| 131401–132000 | 119237–119759 | *ccsA* | *ndhD* | SSC | * |  |
| 140401–141000 | 127827–128402 | *rps15* | *ycf1* | SSC |  | * |
| 144001–144600 | 131283–131868 | *ycf1* | *ycf1* | SSC | * | * |
| 144601–145200 | 131869–132446 | *ycf1* | *ycf1* | SSC | * | * |

**Table 5.** Nucleotide diversity (π) and differences in log-likelihood (LD) scores of the informative windows identified in this study (bold) and previously proposed barcode markers. High values, above the 95^th^ percentile, are indicated in bold.

| Region | Pi | LD |
| --- | --- | --- |
| *ccsA-ndhD* | **0.0258** | 247.12 |
| *matK* | 0.0153 | 136.92 |
| *petD-rpoA* | **0.0246** | 260.79 |
| *petN-trnD* | 0.0228 | **361.07** |
| *psaJ-rps18* | 0.0176 | **309.10** |
| *psbM-trnD* | **0.0292** | **330.41** |
| *psbZ-trnfM* | **0.0246** | **373.97** |
| *rbcL* | 0.0105 | 95.03 |
| *rpl16-rps3* | **0.0345** | 275.89 |
| *rpoB* | 0.0097 | 120.53 |
| *rpoC1* | 0.0103 | 178.60 |
| *rps15-ycf1* | 0.0212 | **284.57** |
| *trnE-trnT* | **0.0241** | **522.51** |
| *trnfM-psaB* | **0.0254** | **375.76** |
| *trnH-psbA* | 0.0126 | **310.47** |
| *trnK-rps16* | 0.0164 | **350.44** |
| *trnL* | 0.0106 | 192.27 |
| *trnT-psbD* | 0.0239 | **291.15** |
| *trnV-atpE* | 0.0128 | **379.77** |
| *ycf1* (1) | **0.0273** | **462.53** |
| *ycf1* (2) | **0.0469** | **313.12** |

**Table 6.**
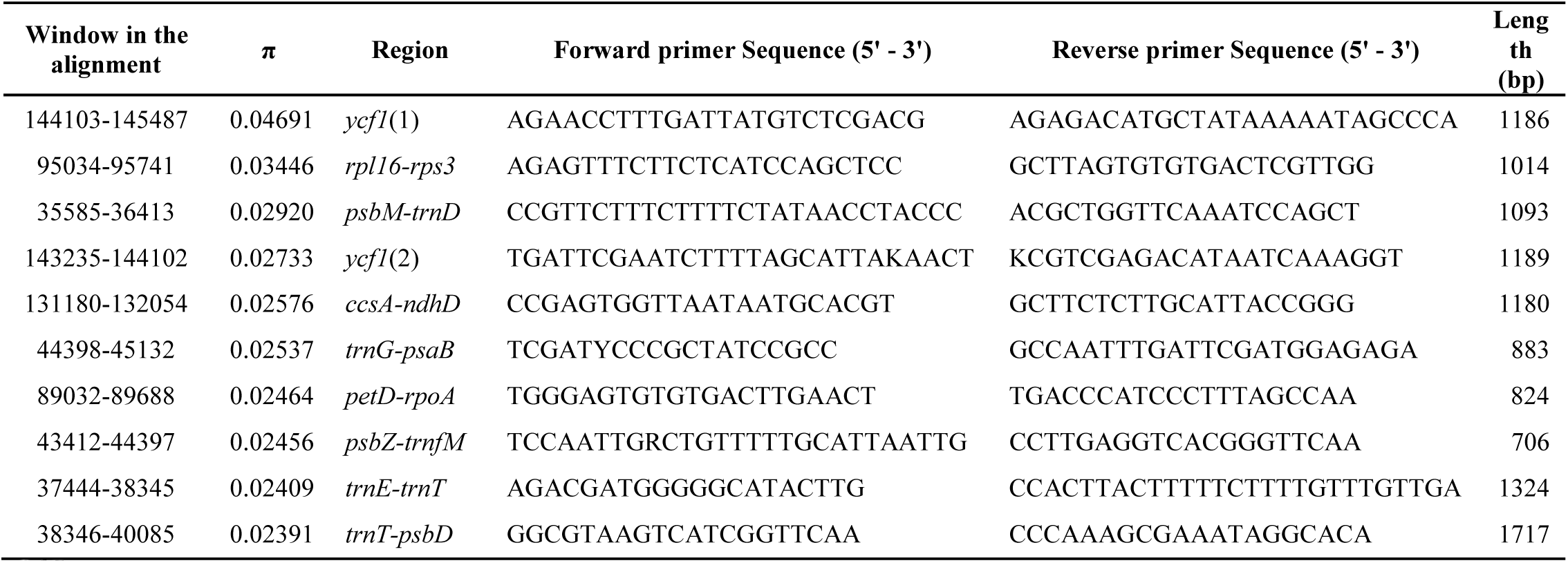
Primer sequences designed to amplify the ten most polymorphic Lecythidaceae plastome regions, as sorted by decreasing nucleotide diversity (π). The product size (length) references the *Bertholletia excelsa* plastome.

**Fig. 2.**
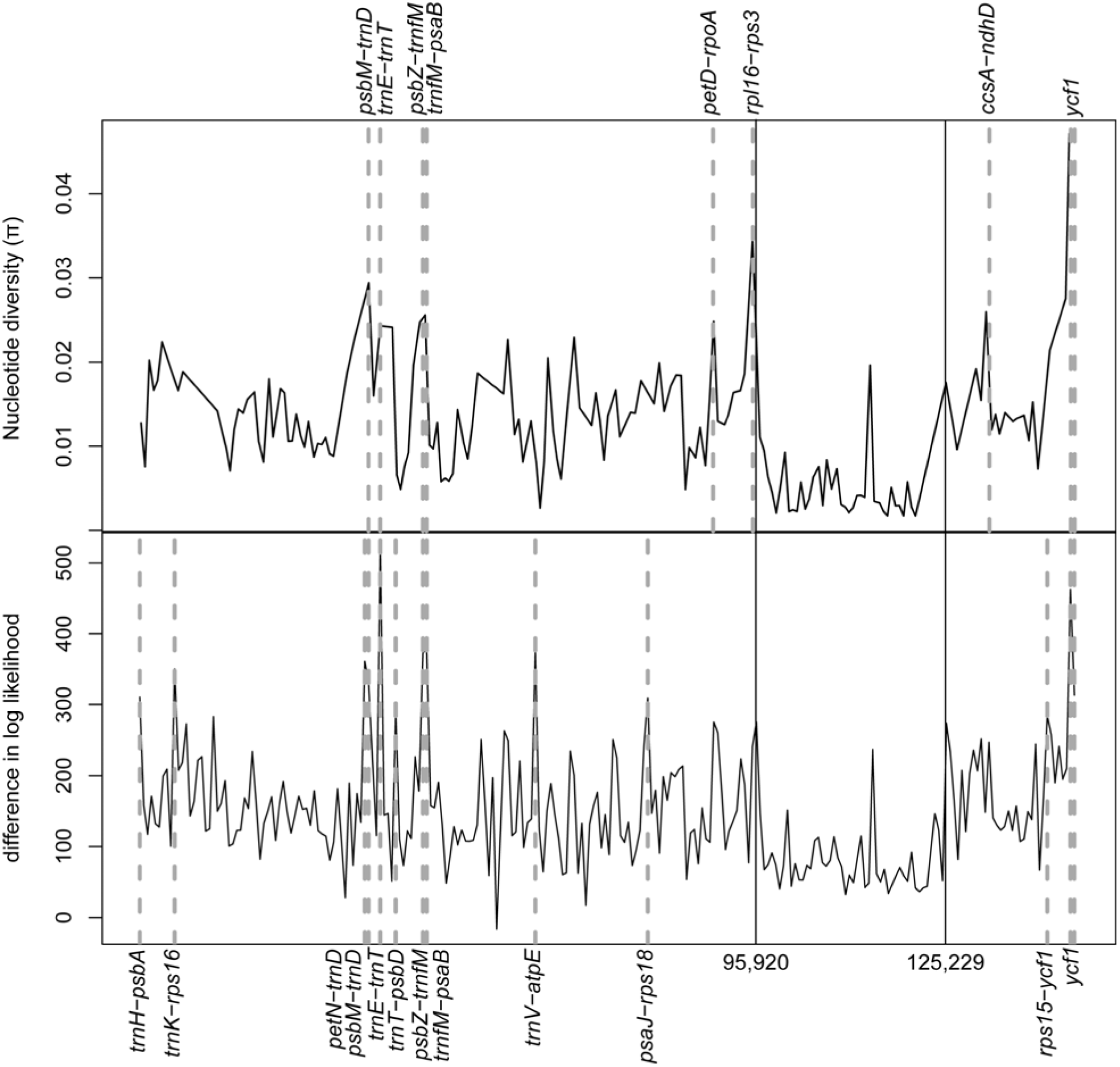
A) Sliding window plot of nucleotide diversity (π) across the alignment of 24 sequenced Lecythidaceae plastomes. B) Alignment site-wise differences in log-likelihood (LD) calculated from the chloroplast topology vs. the average scores of 1000 random trees using a 600-site window. Regions with π and LD above the 95^th^ percentiles are indicated with dashed lines. Continuous vertical lines indicate the boundaries, from left to right, among the large single copy, the inverted repeat, and the small single copy.

***Phylogenetics of the plastomes and the developed markers*—**The ML analysis of the plastome alignment for Lecythidaceae (145,487 sites) yielded a fully resolved phylogeny with high BS for all clades (Fig. 3). Of the genera in which the sampling included multiples species, *Eschweilera* and *Lecythis* were polyphyletic, while *Allantoma, Corythophora, Couratari*, and *Gustavia* were monophyletic *(Bertholletia* is monospecific, and only one species of *Couroupita, Cariniana*, and *Grias* were included in the analysis). The trees obtained from individual markers with high π had an average BS of 73 throughout their nodes, while the trees obtained from two or more concatenated regions had an average BS of 89 (Fig. 4A, Fig. S1). None of the gene trees, single or combined (Fig. S1), recovered the topology obtained using the complete plastome matrix (none of the gene trees obtained a RF=0, Fig. 4B). In general, matrices with concatenated markers (mean RF=6) outperformed single markers (mean RF = 13.8) (Fig. 4B).

**Fig. 3.**
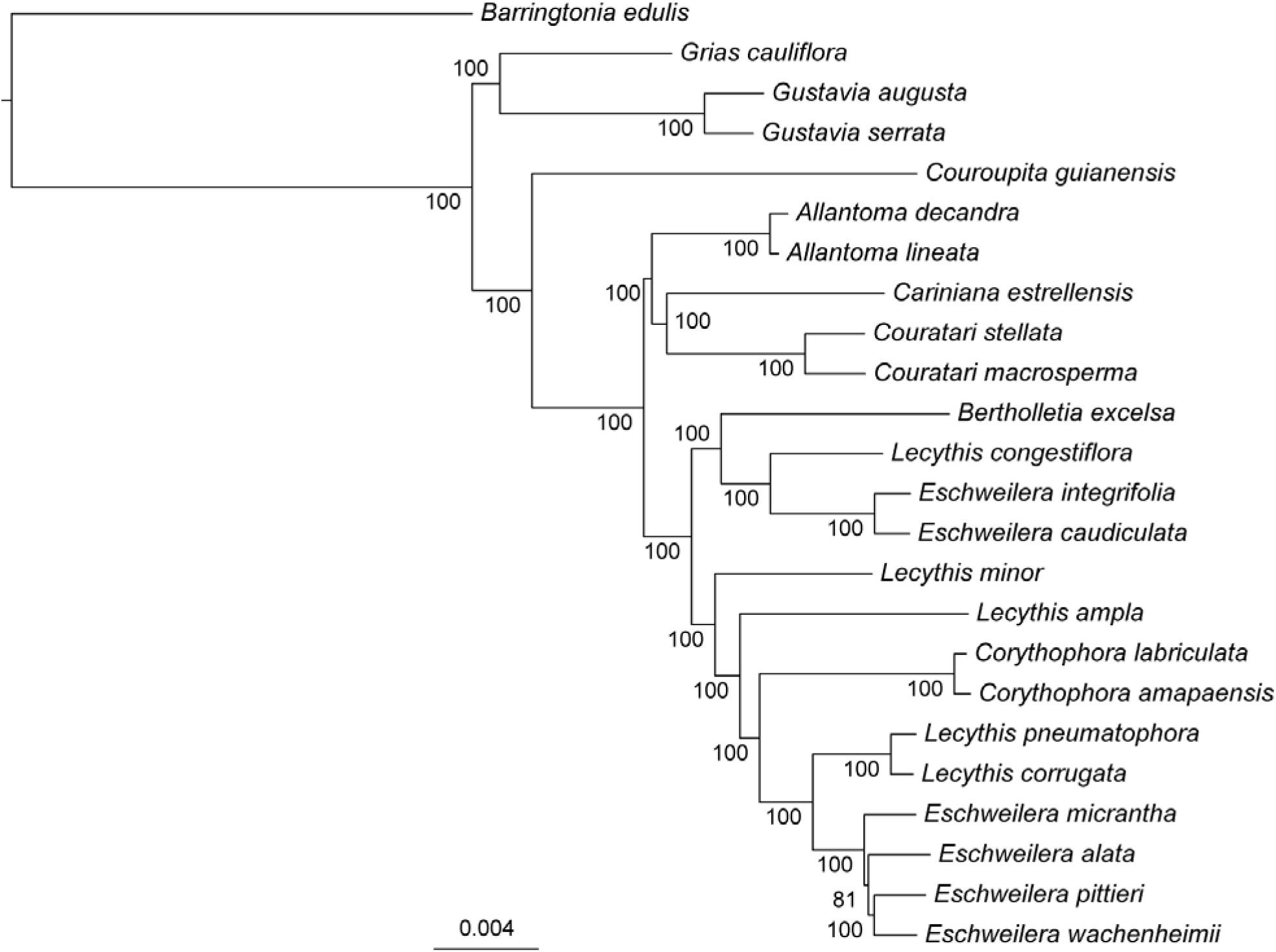
Maximum likelihood phylogeny inferred from plastomes of 23 Neotropical Lecythidaceae. Numbers at nodes indicate bootstrap support.

**Fig. 4.**
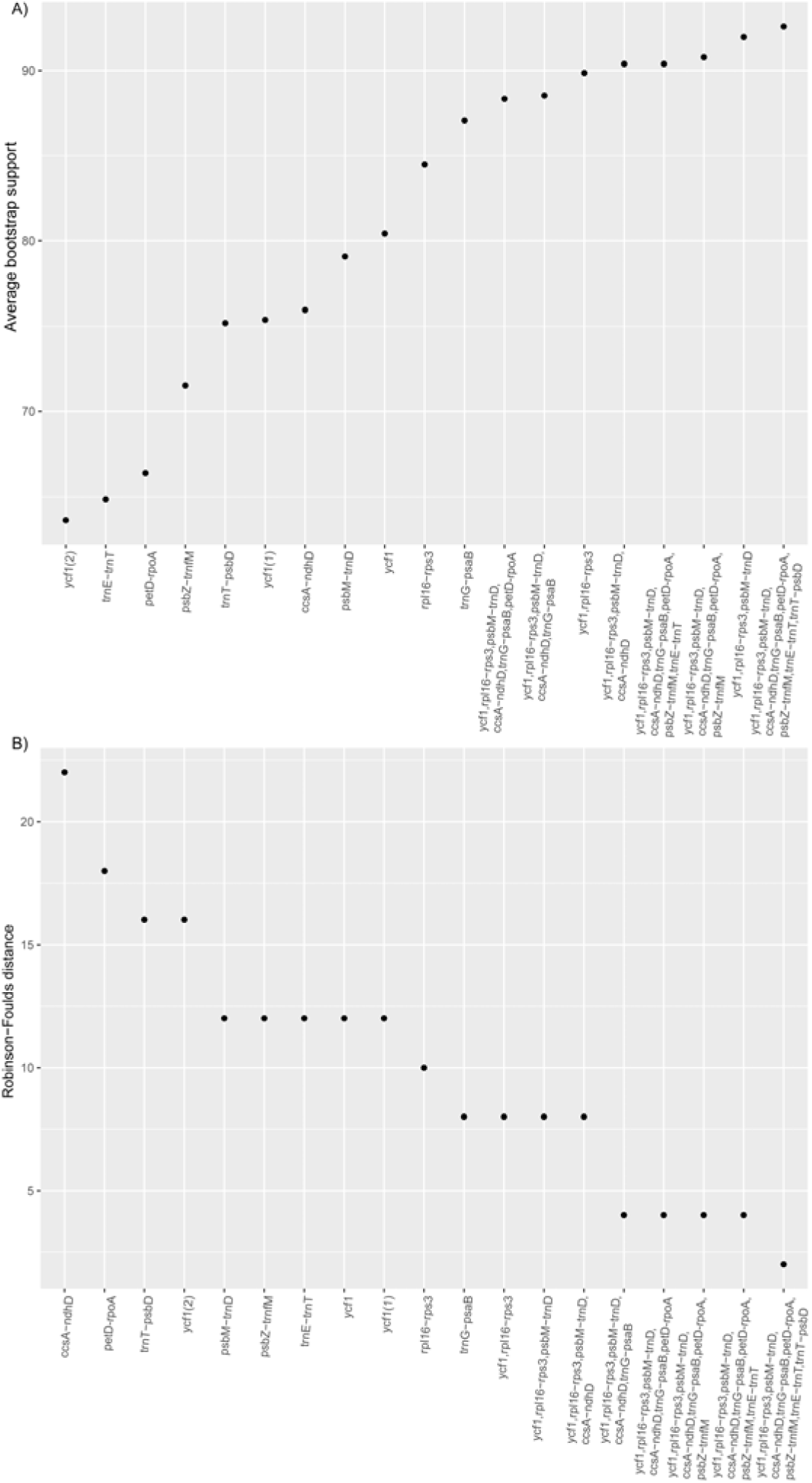
A) Average bootstrap support for trees inferred from matrices of concatenated regions with relatively high nucleotide diversity sorted in ascending order; and B) Robinson-Foulds distance (RF) sorted in descending order. Lower RF distances, which measures the number of different bipartitions from the complete plastome topology, indicate better accuracy.

## DISCUSSION

***Genetic markers from the Lecythidaceae plastome—***We are publishing the first full plastome for Lecythidaceae, including high-depth coverage of the Brazil nut tree, *Bertholletia excelsa*, and 23 draft genomes representing all Lecythoideae genera and a Paleotropical outgroup taxon. We found no significant gene losses or major rearrangements when the plastome of *Bertholletia excelsa* was compared with that of *Camellia sinensis* var. *pubilimba*, a closely related plastome (Theaceae).

We inferred a robust backbone phylogeny for Lecythoideae using the 24 aligned plastomes. All nodes in our topology had 100% bootstrap support except for a node that connects three closely related species of *Eschweilera* (Fig. 3). The topology agreed with previous but weakly supported (<50% BS) Lecythidaceae phylogenies, based on chloroplast and nuclear internal transcribed spacer (ITS) sequences (Mori et al., 2007, Huang et al., 2015), indicating that *Eschweilera* and *Lecythis* are polyphyletic. Although the polyphyly of these two genera is well supported with all available data, some inferred species-level relationships may change with increased taxonomic sampling and the inclusion of nuclear genomic data.

We measured nucleotide diversity (π) and a proxy for phylogenetic signal using a log-likelihood approach (LD) modified from Walker et al. (2017). These calculations helped us to evaluate the performance of specific chloroplast regions as potential phylogenetic markers. The core plant DNA barcodes *matK* and *rbcL* did not exhibit high π or LD in our analysis (Table 5). Of the secondary plastome barcodes mentioned in the literature *(rpoC1, rpoB, trnL, psbA-trnH;* Kress et al, 2005, Lahaye et al., 2008, Hollingsworth 2009, Li et al., 2011) only *psbA-trnH* showed high LD (Table 5) although it did not exhibit exceptionally high values of π. In contrast, the regions *ycf1, rpl16-rps3, psbM-trnD, ccsA-ndhD, trnG-psaB, petD-rpoA, psbZ-trnfM, trnE-trnT*, and *trnT-psbD* displayed the highest values of π and LD and therefore outperformed all the previously proposed plant DNA barcodes.

Phylogenetic trees calculated from concatenated marker sets (based on the π rank) outperformed single regions in terms of support (BS) and accuracy (RF) (Fig. 4). In fact, tree topologies using single markers deviated from the complete plastome tree (mean RF= 13.8). The best performing concatenated matrix contained all 10 regions for which we developed primers. However, the combination of *ycf1* and *rpl16–rps3* produced an average BS ~90 (Fig. 4A) with reasonable accuracy (RF = 4, Fig. 4B); we conclude that these two regions, amplified in three PCRs (Table 6), are promising markers for DNA barcoding, phylogeny, and phylogeography in Lecythidaceae. Although barcoding efficiency in species-rich clades (i.e. *Eschweilera/Lecythis)* might decline with the addition of more samples, *ycf1* and *rpl16-rps3* effectively distinguished between three closely-related species within the *E. parvifolia* clade (see branch lengths in Fig. S1), suggesting that these markers might effectively distinguish between many other closely related species. Our results and conclusions agree with those of Dong et al. (2015) who proposed *ycf1* as a universal barcode for land plants.

The 19 cpSSR markers developed for noncoding portions of the *B. excelsa* plastome provide a useful resource for population genetic studies. Because of their fast stepwise mutation rate relative to SNPs, cpSSRs can also be used for finer grain phylogeographic analyses (e.g. Lemes et al., 2010; Twyford et al., 2013). This may be especially useful for species that exhibit little geographic structuring across parts of their ranges. Because they are maternally transmitted and can be variable within populations, the cpSSRs may also be used to track dispersal of seeds and seedlings relative to the maternal source trees.

Because of their high level of polymorphism and phylogenetic signal content, we anticipate using the cpDNA markers presented here to study the phylogeography of widespread Lecythidaceae species, such as *Couratari guianensis* and *Eschweilera coriacea*, which range from the Amazon basin into Central America.

***Barcoding of tropical trees*—**DNA barcoding of tropical trees has been useful for several applications (Dick and Kress, 2009), including community phylogenetic analyses (Kress et al., 2009), inferring the species identity of the gut content (diet) of herbivores (García-Robledo et al., 2013), and for species identification of seedlings (Gonzalez et al., 2009). The power of DNA barcodes to discriminate among species should be high if the studied species are distantly related; for example, Kress et al. (2009) were able to discriminate 281 of 296 tree and shrub species from Barro Colorado Island (BCI) using standard DNA barcodes, but they were not able to discriminate among some congeneric species in the species-rich genera *Inga* (Fabaceae), *Ficus* (Moraceae), and *Piper* (Piperaceae). Gonzales et al. (2009) encountered similar challenges with *Eschweilera* species in their study of trees and seedlings in Paracou, French Guiana. The latter study tested a wide range of putative DNA barcode regions *(rbcLa, rpoC1, rpoB, matK, trnL, psbA-trnH*, ITS) but did not include the markers presented in this paper.

***Limitations of plastome markers for phylogeny and species ID*—**These newly-identified plastome markers are not free of limitations. First, plastome-based phylogenies should be interpreted with caution, as they can disagree with nuclear markers and species trees due to introgression and or lineage sorting issues (Rieseberg and Soltis, 1997; Sun et al., 2015; Vargas et al., 2017). These same processes limit the cpDNA for species identification. For example, cpDNA haplotypes of *Nothofagus, Eucalyptus, Quercus, Betula*, and *Acer* were more strongly determined by geographic location than by species-identity due to the occurrence of localized introgression within these groups (Petit et al., 1993; Palme et al., 2004; Saeki et al., 2011; Premoli et al., 2012; Nevill et al., 2014; Thomson et al., 2015). The occurrence of haplotype sharing in closely-related Lecythidacae species has, to date, not been examined at a large scale and it is therefore not possible to conclude to what extent introgression or incomplete lineage sorting might affect this group. We suggest that future studies utilizing cpDNA barcodes for Neotropical Lecythidaceae test species from several shared geographic localities to examine to what extent haplotypes tend to be shared among species at the same localities. Nuclear DNA markers may also be used to examine phylogenetic incongruence and identify cases where introgression might have occurred.

## ACKNOWLEDGEMENTS

National Science Foundation (DEB 1240869 and FESD Type I 1338694 to CD) and the University of Michigan (Associate Professor Award to CWD) provided financial support for this work. We would like to thank Scott Mori, Gregory Stull, Caroline Parins-Fukuchi, Joseph Walker, and three anonymous reviewers for their useful comments; and Scott Mori and the New York Botanical Garden for providing access to curated DNA samples of Lecythidaceae.

## DATA ACCESSIBILITY

DNA sequences: Genbank accessions MF359935–MF359958, BioProject SUB2740669

Plastome alignment, gene alignments, trees, and R code: https://bitbucket.org/oscarvargash/lecythidaceae_plastomes

## SUPPORTING INFORMATION

Fig. S1. Phylograms obtained from single and combined markers with high nucleotide diversity. Numbers at nodes indicate bootstrap support.

Appendix 1. Lecythidaceae species sequenced with their voucher, assembly information, and GenBank accession number. All voucher specimens are deposited at the New York Botanical Garden Herbarium (NY).

